# Structured connectivity for structured sequential computations

**DOI:** 10.64898/2026.08.03.742515

**Authors:** Kalel L. Rossi, Aneta Koseska

## Abstract

Cortical computations emerge from the coordinated activity of recurrent neural circuits and are often described through population dynamics and neuronal selectivity. However, how the backbone of the computations - the circuit connectivity - gives rise to these dynamics and organizes the computations, remains largely unknown. Mixed selectivity, which can arise without structured connectivity, has been proposed as a substrate for flexible computation, but alone it does not account particularly for sequential computations, where distinct selectivities have been observed. Here, we investigate this problem using the delayed match-to-category (DMC) task, which requires categorization, memory maintenance, comparison, and decision-making, and in which pure, mixed, and time-varying selectivity have been observed. Using explicit connectivity models, we identify circuit motifs that generate these selectivity types and interconnect them to implement the computations required for behavior. Within these motifs, mixed selectivity neurons implement comparison via localized, distributed, or hybrid representations, whereas time-varying selectivity neurons gate the activation and timing of output. Recurrent networks trained from random initial connectivity develop similar motifs, demonstrating that learning leads to structured connectivity to support computations. The framework generalizes to extended task variants, revealing reusable circuit principles that mechanistically link connectivity, neuronal selectivity, and sequential computation.

## I. INTRODUCTION

Cortical computations occur sequentially in many behavioral paradigms, with the population activity implementing consecutive computational stages. In this way, the output from one stage serves as input to the next [1–5]. Such sequential transformations underlie diverse cognitive functions, including sensory processing, working memory and decision-making, and have been extensively characterized in recurrent neural circuits [1–5]. However, how circuit connectivity organizes these successive computational stages remains poorly understood. A simple yet powerful paradigm for investigating this gap is the delayed match-to-category (DMC) task, in which subjects must transform sensory stimuli into abstract categories, maintain category information across a delay, compare the stored category with a subsequent stimulus, and generate an appropriate match or non-match decision (Fig. 1A) [6–8]. This sequential organization provides a natural framework to examine how circuit structure supports the transformation of information between distinct computational stages.

**Figure 1.**
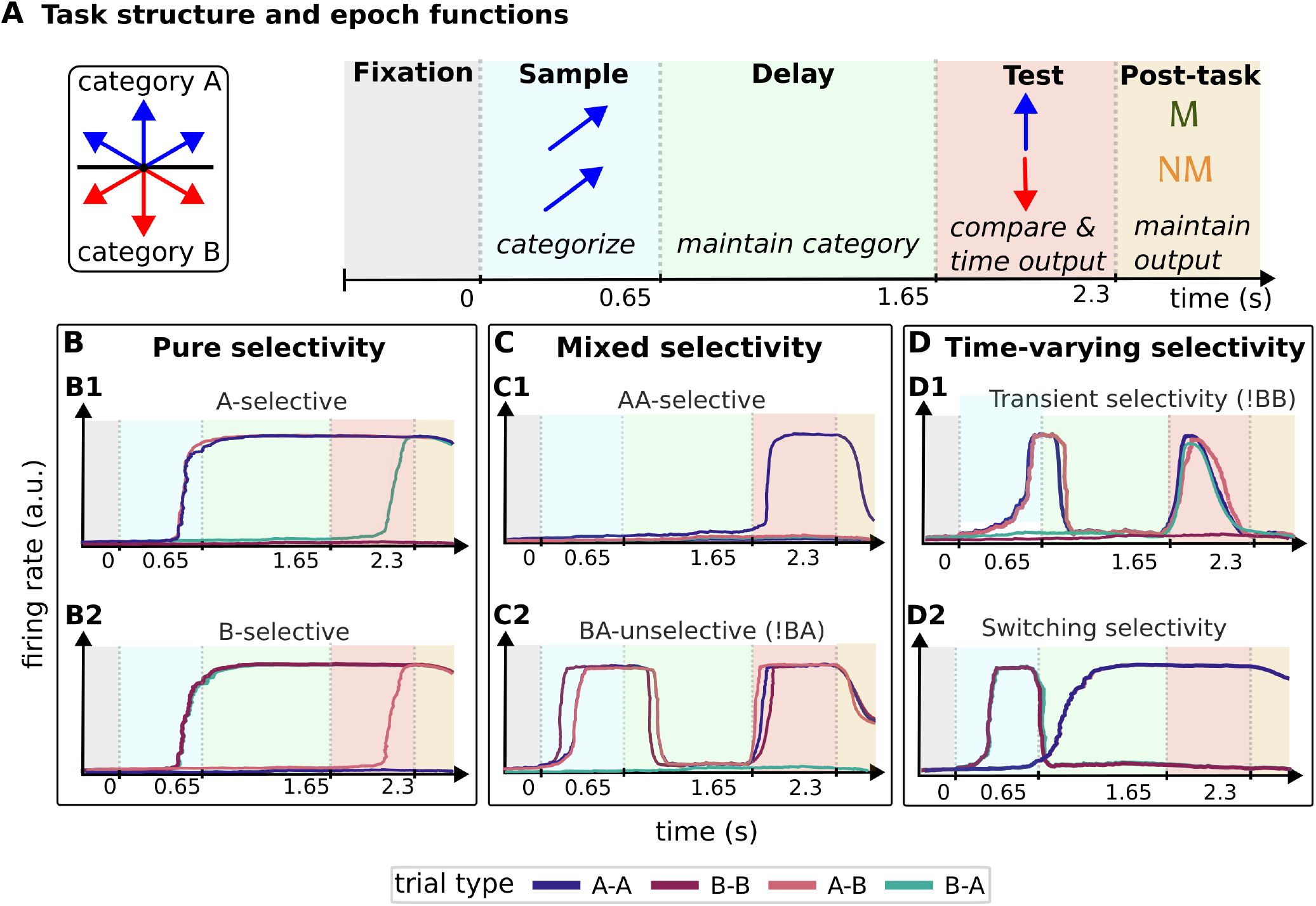
Structure of delayed match-to-category (DMC) task and illustrative examples of experimentally observed neural traces. (A) In the DMC task, primates view two sequentially presented visual stimuli consisting of coherently moving dots, separated by a delay period [6–8]. Stimuli are represented by their motion direction (angle), and subjects compare the categories of the two stimuli to report a match (AA, BB) or non-match (AB, BA) decision [3]. (B) Schematic neuronal activity traces representing experimental recordings from prefrontal cortex and lateral intraparietal cortex, revealing diverse neuronal response dynamics, including pure selectivity (PS, responses selective for individual categories), mixed selectivity (MS, responses to combinations of task variables) [6, 9], and time-varying selectivity (TVS, response properties that change across task epochs [3])

Recordings from primate prefrontal and parietal cortex during the DMC task reveal strikingly heterogeneous neuronal dynamics (Fig. 1B-D). Individual neurons exhibit diverse response profiles: some neurons respond selectively and encode category identity throughout the task (pure selectivity, PS), others respond to combinations of task variables (mixed selectivity, MS) [3, 4, 6], while many change their response properties across task epochs (time-varying selectivity, TVS) [3, 10, 11]. Similar mixed-selective representations have also been reported in mouse parietal cortex during delayed match-to-sample tasks, suggesting that such MS is a general feature of cortical computations rather than a specialization of primate prefrontal cortex [9]. Existing theories have provided important insights into the computational role of neuronal selectivity. Random recurrent networks naturally generate mixed-selective neurons, producing highdimensional representations that support flexible linear decoding for computations [12–16]. However, this framework does not readily explain the emergence of the stable category-selective representations required for the DMC task and consistently observed in cortical recordings [3]. To account for these observations, modeling studies have proposed that category-selective specialized populations provide stable representations and project onto mixed-selective neurons responsible for downstream computations [3, 17]. Although these models capture key aspects of recorded neuronal activity, they leave unanswered a more fundamental question: how does circuit connectivity organizes these distinct neuronal representations into the sequential of computations required for behavior? Consequently, a mechanistic framework linking circuit connectivity, neuronal selectivity, and the temporal organization of computation is still lacking.

Here, we develop minimal circuit models that explicitly reveal how recurrent connectivity organizes the successive computations underlying the DMC task. The circuits first transform sensory inputs into stable categorical representations maintained throughout the delay and then sends these representations to structured comparison circuits that inform the match or non-match decision, which is also controlled by gating neurons. This circuit organization naturally gives rise to pure-selective, mixed-selective, and time-varying neuronal responses, assigning each selectivity type a distinct computational role. We further show that stimuli comparison can be implemented through multiple connectivity architectures spanning localized and distributed representations [18], providing an integrated explanation for the diversity of population activity observed experimentally and in previous modeling studies [3]. Rather than arising as a by-product of recurrent dynamics, time-varying selectivity emerges as a functional mechanism for coordinating the timing of decision formation. Finally, we show that recurrent neural networks trained from random initial connectivity consistently converge on connectivity motifs closely resembling those derived in the minimal models. This convergence suggests that learning naturally discovers structured circuit solutions that organize sequential computations through specialized connectivity, rather than relying on arbitrary distributed dynamics, reminiscent of findings showing structured activity encoding specific behavioral variables in the rat orbitofrontal cortex in a decision-variable integration task [19–21].

Together, our results establish a mechanistic link between circuit structure, neural dynamics, and computation, integrating experimental observations into a concrete and detailed description of how structured connectivity gives rise to the sequential neural dynamics that underlie flexible behavior.

## II. RESULTS

### A. Category encoding and comparison are implemented by distinct circuit modules

We hypothesized that successful DMC performance relies on two core computational operations: category encoding and category comparison. Category information must first be extracted from the sample stimulus and maintained throughout the delay period, after which this internal representation must be integrated with the category of the test stimulus to generate a match or non-match decision (Fig. 1A). This decomposition suggests that behavior may be supported by distinct but functionally coupled circuit components.

To identify the minimal circuit architectures capable of implementing these computations, we constructed neuronal circuits constrained not only to solve the task but also to reproduce the diversity of neuronal dynamics observed experimentally. Rather than attempting to capture the full complexity of cortical networks, these models were designed to identify the minimal architectural principles sufficient to generate pure selectivity (PS), mixed selectivity (MS), and time-varying selectivity (TVS) while producing accurate behavior. In all models, direction-selective stimulus input neurons projected excitatory and inhibitory connections onto neuronal circuit with continuous-time rate dynamics (see Methods; Fig. 2 A, D, G). The output layer consisted of two decision neurons encoding match (M) and non-match (NM) responses. Across architectures, category encoding was implemented through PS neurons that selectively represented individual categories and maintained these representations throughout the delay through recurrent self-excitation, forming a dedicated category-encoding module. Comparison was implemented by downstream mixed-selectivity (MS) populations that integrated the maintained category representation with information about the test stimulus. Consistent with neural coding theory [18], we found that this computation was not associated with a single canonical architecture, but instead emerged through multiple circuit motifs that generated distinct forms of mixed selectivity, ranging from localized to distributed representations.

**Figure 2.**
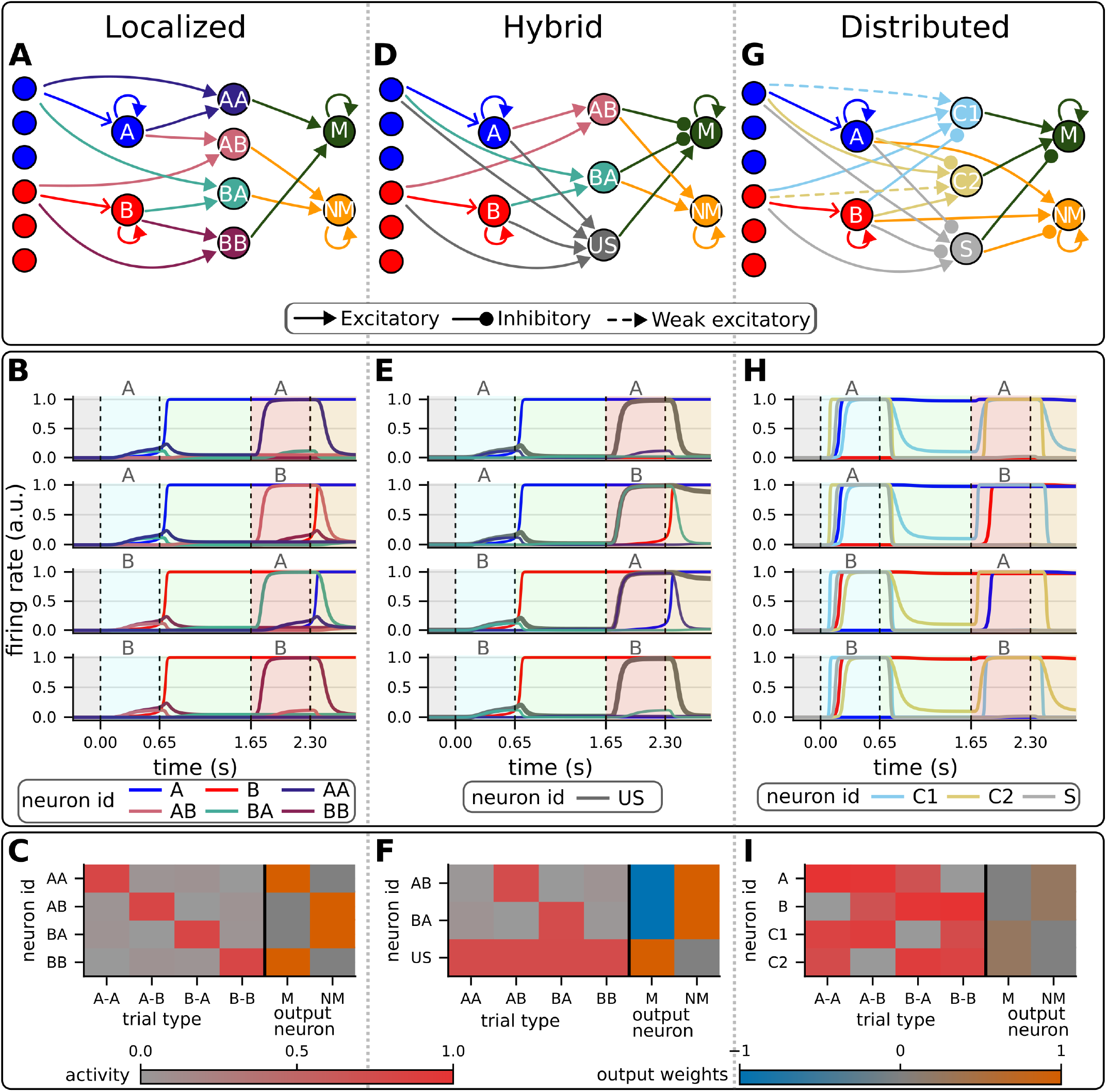
Minimal circuit architectures implementing DMC computations. Minimal circuits were constructed to perform the DMC task while reproducing experimentally observed neuronal dynamics (Fig.1B-D). For visualization, only six input neurons are shown, each tuned to a specific motion direction; neurons tuned to category A are colored blue and category B are red. For clarity, connections from only one input neuron are displayed; the full networks contain the same connectivity structure replicated across all input units. (A–C) Localized comparison motif. (A) Circuit architecture in which individual mixed-selective neurons encode specific sample–test category combinations. (B) Respective network activity across trial types, showing the emergence of pure- and mixed-selective responses. (C) Corresponding activity of neurons contributing to the decision output, averaged across the decoding period, together with their output weights. (D) Hybrid comparison motif, combining localized and distributed representations, with an unspecific-selectivity population contributing to the decision computation. (E,F) Corresponding neuronal responses and decoding strategy. (G) Fully distributed comparison motif, in which individual neurons do not encode trial identity; instead, match and non-match decisions emerge from the collective activity of multiple neurons (F,I).

In the first motif, we identified a localized solution in which the comparison computation emerged through mixed-selective neurons that explicitly represented specific combinations of sample and test categories (AA, AB, BA, or BB) (Fig. 2A, B). These neurons functioned as conjunction detectors, analogous to AND gates, receiving convergent inputs from category-selective PS neurons and test-category representations. Because each trial type was represented by a dedicated neuronal population, the decision variable was explicitly encoded at the single-neuron level, with AA and BB populations driving match responses and AB and BA populations driving non-match responses (Fig. 2C; Supp. Materials and Supp. Fig. 1A). In a second motif, we identified a hybrid solution in which the comparison computation emerged through partially distributed representations (Fig. 2D-F). In this architecture, an unspecific (US) neuron provided a common drive to the match output, while inhibitory inputs from AB and BA neurons selectively suppressed this response during non-match trials. As a result, match and non-match information was not localized to individual neurons but emerged from the coordinated activity of multiple neurons, producing a distributed two-dimensional representation of trial identity (Supp. Fig. 1B). Finally, we identified a fully distributed solution in which comparison emerged without neurons explicitly encoding individual trial types (Fig. 2G-I). Instead, neurons active across match trials collectively drove the match output, whereas neurons preferentially active during nonmatch trials drove the non-match output. Trial information was therefore compressed onto a population-level decision axis separating match and non-match responses and could only be recovered from the collective activity of the network (Supp. Fig. 1C). Interestingly, the same neuronal activity patterns also supported localized decoding when inhibitory connections onto the output layer were introduced (Supp. Materials and Supp. Fig. 2), demonstrating that the computational role of a neuronal population depends not only on its activity structure but also on the connectivity through which this activity is read out. Notably, in this circuit realization, the activity profiles of neurons C1 and C2 resemble mixed-selective responses (Fig. 1C(C2)) observed in lateral intraparietal area during similar tasks [3], suggesting that these minimal motifs capture circuit principles underlying experimentally observed cortical dynamics.

Despite their distinct representations, all motifs implement the same underlying computational decomposition: maintaining a stable category representation and combining it with sensory evidence to generate a decision. In each architecture, PS neurons provide a persistent representation of category identity, which is transformed by downstream interactions with test-stimulus information to produce the comparison required for behavior. Thus, the same sequence of computations can be realized through different connectivity organizations, ranging from localized to distributed representations. These minimal circuits reveal that heterogeneous neuronal responses need not reflect arbitrary variability, but can instead emerge as distinct architectural solutions to a shared computational objective. Pure and mixed selectivity therefore represent different circuit-level implementations of how information is maintained, transformed, and routed to support flexible behavior.

### B. Time-varying selectivity gates the timing of decision

The minimal circuits in Fig. 2 establish how category encoding and comparison can be implemented through distinct circuit architectures. However, successful DMC performance requires not only the correct computations, but also their precise temporal coordination. Because the information available to the circuit changes across task epochs, decision-related activity must remain suppressed during stimulus encoding and delay periods and become engaged only when the relevant sensory evidence is available. This suggests that neural circuits require mechanisms that regulate the timing of information flow between computational stages.

To identify how such temporal control can emerge, we extended the minimal architectures by incorporating neurons (*S, T*, R^0^, and R_p_) that regulate the timing of computation (Fig. 3A). Sample-selective (*S*) neurons suppress decision-related populations during stimulus presentation, preventing premature activation of the output layer (Fig. 3B). Conversely, test-selective (*T*) neurons become activated after test onset and facilitate the engagement of comparison and decision circuits once sufficient information is available. A distinct mechanism is provided by ramping (R_p_) neurons, which gradually increase their activity throughout the delay period and peak around test onset, providing an internal timing signal for when comparison should be initiated. In our model, this ramping activity emerges through transient dynamics generated by a ghost channel [22]: a transiently active neuron (R^0^) gradually drives the activation of R_p_, producing a temporally evolving signal analogous to neural integrator mechanisms [23]. Although ramping and time-dependent activity patterns are widely observed in frontal and parietal cortex during delayed decision and working-memory tasks [9], the circuit mechanisms that generate these dynamics and their functional contribution to sequential computation have remained unclear. Our model however suggests that these temporal signals can provide a mechanistic substrate for coordinating when distinct computations are engaged within a circuit.

**Figure 3.**
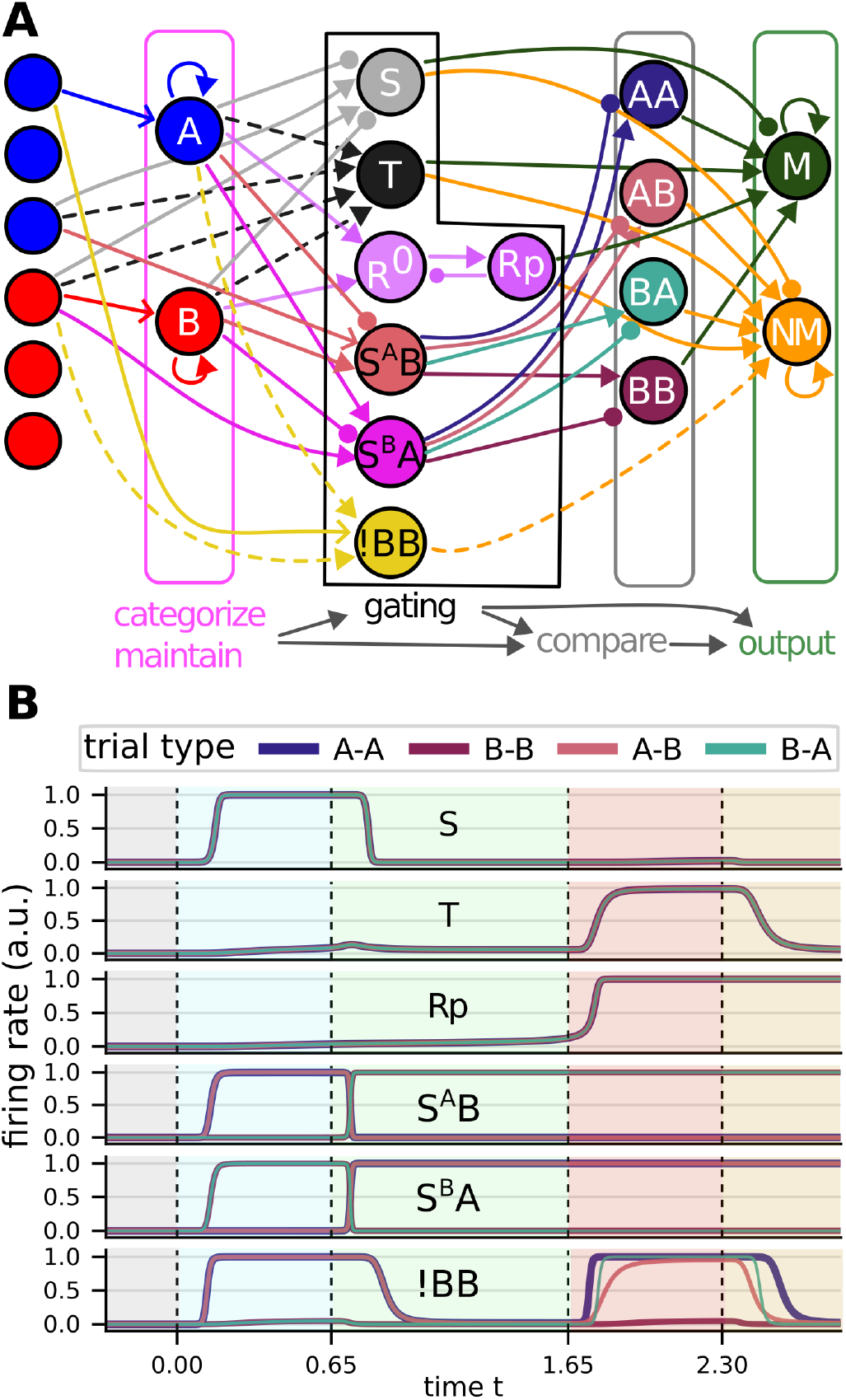
Augmented circuit architectures generating time-varying selectivity. (A) Connectivity of an augmented network incorporating auxiliary neurons that generate time-dependent selectivity profiles. (B) Neuronal activity across trial types. Epoch-selective neurons provide temporal control of computation: sample (S) neurons regulate activity during sample encoding, whereas test (T) neurons facilitate decision generation at the appropriate time. T neurons excite the output layer, while S neurons inhibit output during the sample period to prevent premature responses. Ramping neurons (*R*^0^, *R*_*p*_) provide additional temporal modulation of the decision process. Multiplexing neurons combine multiple computational roles across trial types; for example, S^B^A functions as a sample-selective neuron for BB and BA trials and as a category-selective neuron for AA and AB trials, whereas S^A^B exhibits the complementary response pattern. The !BB neuron performs the same computational role as the A-selective population in the fully distributed motif (Fig. 2G), but without maintaining delay-period activity.

Importantly, temporal control does not require category-unselective gating neurons. Equivalent regulation of computations can be achieved by replacing a single unselective gating neuron with a pair of category-selective neurons that share the same temporal profile but respond selectively to different categories. While this preserves circuit function, the activity of each individual neuron now exhibits category-specific responses restricted to a particular task epoch, producing response patterns resembling experimentally observed time-varying selectivity [9]. Motivated by time-varying responses reported in prefrontal cortex [3, 6, 7], we next explored architectures in which neurons dynamically change their computational role across task epochs. Neurons S_A_B and S_B_A multiplex sample and category representations by acting as sample-selective neurons during one epoch and category-selective neurons during another (Fig. 3B). Their changing activity profiles generate switching selectivity [3], reflecting transitions in functional role across the trial. This multiplexing allows a single population to participate in multiple computations but introduces constraints imposed by fixed connectivity: when these neurons become category-selective, they may also provide unnecessary drive to comparison circuits during earlier epochs. Thus, temporal multiplexing provides computational flexibility at the cost of additional requirements for connectivity regulation. A related example is provided by the !BB neuron, inspired by time-varying responses observed in LIP (Fig. 1D (D1), Fig. 3). In the distributed comparison motif (Fig. 2G), this neuron performs the computational role of the Aselective population but lacks delay-period activity, preventing premature activation of the output layer. This illustrates how time-varying selectivity can arise from architectural constraints that separate when information is represented from how it is used.

Together, these results provide a mechanistic account for the functional role of time-varying selectivity. Rather than reflecting passive fluctuations in heterogeneous neural activity, TVS can emerge as a circuit strategy for coordinating the engagement of different computations and enabling neurons to participate flexibly across task stages. Thus, heterogeneous and time-varying selectivity reflect not only how information is represented, but also how circuit architectures organize the temporal flow of computations underlying behavior.

### C. Trained networks spontaneously develop structured computational motifs

The minimal models above identify circuit architectures sufficient to implement the computations required by the DMC task. We next asked whether these architectural principles emerge spontaneously in recurrent neural networks (RNNs) trained from random initial conditions. Following previous modeling work [3], we trained continuous-time recurrent neural networks consisting of 32 direction-selective input neurons with von Mises tuning, a randomly initialized recurrent population, and two output neurons encoding match (M) and non-match (NM) decisions using backpropagation through time [24] (see Methods, Sec. VI B). We then examined whether training organizes connectivity into the computational motifs predicted by the minimal circuits.

Despite random initialization, trained networks converged onto structured circuit organizations (Fig. 4A; Supp. Fig. 3A, B). Pure-selective populations emerged for each category and formed stable recurrent assemblies characterized by strong within-category excitation (Fig. 4B; Supp. Fig. 3C). These populations maintained category information throughout the delay period, implementing an encoding stage analogous to that identified in the minimal circuits (Fig. 2A, D, G). Downstream of this encoding populations, the trained networks developed heterogeneous comparison populations that combined stored category information with the test stimulus. Some neurons exhibited localized mixed selectivity, responding to specific sample-test combinations and preferentially driving non-match decisions, whereas others encoded distributed comparison signals that were only recoverable at the population level (Fig. 4C; Supp. Fig. 3D). In addition, the networks also developed epoch-selective neurons that became active selectively during the test period and strongly influenced the decision output (Fig. 4D; Supp. Fig. 3E). Thus, rather than converging onto a single representational strategy, learning produced a hybrid architecture combining localized, distributed, and temporally gated computations within the same circuit.

**Figure 4.**
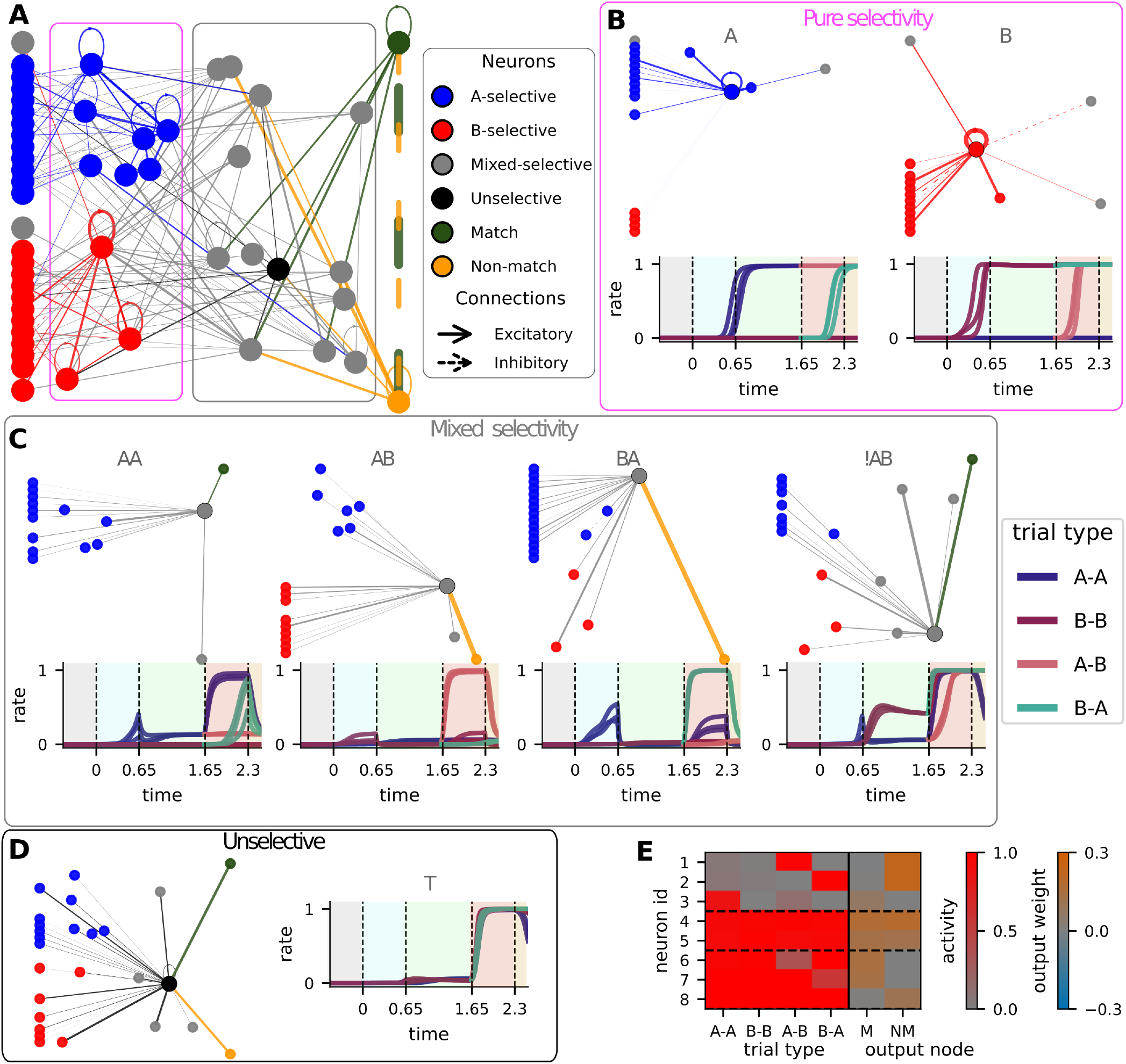
Connectivity and selectivity organization in trained network. (A) Connectivity structure of a representative trained recurrent neural network. Nodes are colored and positioned according to their selectivity class, identified from their activity profiles, and edge widths indicate the magnitude of synaptic weights. For visualization, weak connections and nodes lacking strong connections are omitted. (B–D) Connectivity and activity profiles of representative neurons from the trained network. Top, incoming and outgoing connections of selected neurons; bottom, neuronal responses across all stimulus combinations, grouped by trial type. (E) Activity and output connectivity of neurons with the strongest influence on decision generation. Left, mean firing rates during the decoding period for each input combination within a trial type (gray-to-red scale). Right, corresponding output weights onto match and non-match decision neurons (blue-to-brown scale).

To understand how these representations contribute to behavior, we examined the decoding period, when recurrent activity is transformed into categorical decisions through the output layer. Neurons with the strongest output projections revealed a functional division of labor: localized mixed-selective populations contributed predominantly to non-match decisions, whereas match decisions relied more heavily on distributed and category-independent comparison signals (Fig. 4E; Supp. Fig. 3F). Consistent with the augmented minimal circuits (Fig. 2), trained networks also developed neurons with epoch-dependent response profiles resembling experimentally observed time-varying selectivity (Fig. 4D, Supp. Fig. 3E). These results demonstrate that learning integrates multiple computational motifs within a single circuit rather than selecting a unique architectural solution.

Overall, RNNs trained on the DMC task converge from random initial conditions onto structured circuit architectures composed of stable category-encoding populations, heterogeneous comparison circuits, and temporal control mechanisms. This emergent organization supports the idea that the motifs identified in the minimal models reflect fundamental computational constraints imposed by the task, rather than manually specified design choices. At the population level, these architectures generate trajectories that first separate according to category identity, stabilize during the delay period, and subsequently transition toward match or non-match decisions after test onset (Supp. Fig. 5), in agreement with previous findings [3]. Stable states support persistent category memory, whereas transient dynamics coordinate transitions between successive computational stages. Importantly, similar population trajectories arise from circuits implementing distinct comparison architectures, indicating that neural dynamics alone are insufficient to identify the underlying computation. Instead, neuronal function emerges from the interaction between activity patterns and the connectivity that transforms those patterns into task-relevant computations, supporting the architectural principles identified by the minimal models.

**Figure 5.**
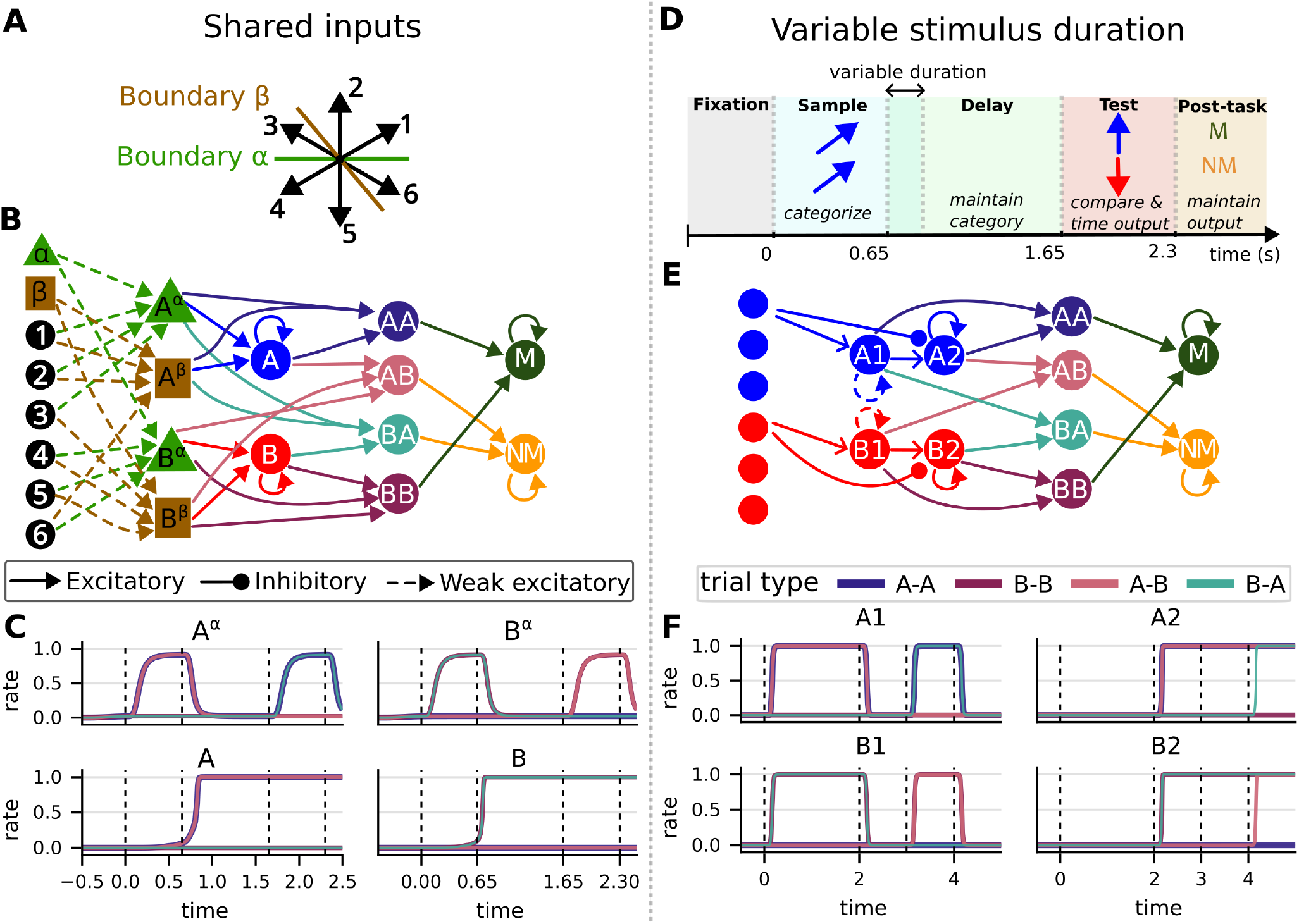
Minimal circuit architectures for extended DMC tasks. (A) A context cue, presented throughout the trial, specifies the category boundary used to classify motion directions. (B) Circuit architecture for a context-dependent DMC task. Context-dependent categorization is implemented through an intermediate layer that combines cue and stimulus information before category encoding. (C) Representative activity traces for the context-specific category populations under the *α* context; analogous responses are observed for the *β* context. (D) Schematic of the DMC task with variable sample / delay durations. (E) Circuit architecture in which temporal flexibility is achieved by separating category encoding into populations active during the sample (A1/B1) and test (A2/B2) epochs, allowing the network to maintain correct computations despite variable trial timing. (F) Representative neuronal activity illustrating the sequential engagement of these populations across the task.

### D. Structured motifs predict circuit adaptations to new task demands

A key advantage of identifying minimal computational motifs is that they provide a predictive framework for how circuit architectures could adapt when behavioral demands change. Rather than representing solutions specific to the standard DMC task, the motifs define reusable computational components whose organization can be modified to accommodate new task requirements. We therefore asked how the proposed architectures should be extended to solve more complex variants of the DMC task.

We first considered a context-dependent DMC task, in which category boundaries depend on an external contextual cue (Fig. 5A). This task introduces an additional computational requirement: sensory information must be transformed into context-dependent category representations before they can be maintained and compared. This requirement can be accommodated in the minimal models by augmenting the encoding stage with a context-dependent routing module that combines sensory and contextual information while preserving the downstream comparison and decision circuitry (Fig. 5B, C). Consistent with previous experimental observations of context-dependent population dynamics [3] and neurons jointly encoding category and task rule [25], this architecture predicts that neural trajectories should initially diverge according to context before converging onto shared comparison and decision mechanisms. We next examined task variants with variable stimulus and delay durations (Fig. 5D), which require category memory to remain stable despite unpredictable timing. Unlike the original minimal circuits (Fig. 2), which exploit a fixed temporal relationship between stimulus offset and memory maintenance, this task requires separating transient sensory encoding from persistent category representations. In the minimal models, this can be achieved by introducing sequential category-selective populations: an initial stimulus-driven population (A_1_/B_1_) transfers activity to a recurrent memory population (A_2_/B_2_) that maintains category information until the comparison stage is engaged (Fig. 5E, F). This additional recurrent layer preserves the computational organization of the circuit while decoupling memory from stimulus duration.

Together, these extensions demonstrate that the proposed motifs constitute a general architectural framework rather than task-specific solutions. New behavioral demands are accommodated by augmenting individual computational stages while preserving the core organization linking category encoding, comparison, and temporal control. This modularity suggests that recurrent circuits are organized around reusable computational principles that can be recombined to support increasingly flexible behavior, providing experimentally testable predictions for how circuit architecture should evolve as cognitive demands increase.

## III. DISCUSSIONS

Flexible behavior requires neural circuits to transform information through a sequence of computations, in which the output of one operation becomes the input of the next. Using the delayed match-to-category task as a canonical example of sequential computation, we show that circuits can decompose the task into three core computations category encoding, comparison, and temporal control and that each can be implemented by identifiable circuit motifs that generate distinct forms of neuronal selectivity. Rather than treating heterogeneous neural responses as a by-product of recurrent dynamics alone, our work establishes a mechanistic relationship between circuit architecture, neuronal dynamics, and computation, providing an explicit structure–dynamics–function relationship for understanding cognitive circuits.

A central result of this work is that the same computation can be realized through multiple circuit organizations. Our minimal models demonstrate that comparison can emerge through localized, distributed, or hybrid architectures, each producing different patterns of mixed selectivity [18]. These findings suggest a reconciliation between two views of cortical computation that have often been considered separately. On one hand, mixed selectivity has been proposed to arise from random, highdimensional recurrent dynamics that enable flexible computation [12, 15]. On the other, experimental evidence also suggests that cortical circuits contain specialized populations encoding specific task variables [19–21]. Our results indicate that localized and distributed representations can both be reached from training to solve a task, and can even co-exist in a given system. Therefore, it would be expected that individuals can achieve similar performances even with distinct neural selectivity structures [26].

An important implication of our results is that neural population dynamics alone do not uniquely determine the underlying computation. In the trained networks and the minimal models, similar trajectories emerged from circuits implementing distinct comparison mechanisms, while still reproducing key experimental features, including stable category representations followed by decision-related transitions. Consistent with recent work showing that neural manifolds do not uniquely determine the underlying computations [27], our results demonstrate that neuronal dynamics alone are insufficient to infer circuit function. Computation depends jointly on activity and on the connectivity that decodes the representations. Understanding cortical computation therefore requires integrating analyses of neural dynamics with circuit architecture rather than considering population trajectories in isolation. This perspective also provides a mechanistic interpretation of time-varying selectivity, which has been widely observed in prefrontal and parietal cortex during delayed decision-making tasks [3, 10, 11]. Whereas previous studies have viewed TVS primarily as a consequence of evolving network states or changing population representations [3], our models suggest a functional role: time-varying selectivity emerges from structured recurrent connectivity to sequentially gate information flow and coordinate the timing of decision formation.

The minimal models introduced here therefore serve as mechanistic building blocks for understanding cognitive computation. Rather than reproducing neuronal activity alone, they identify the essential circuit motifs that implement successive computations, thereby offering a conceptual framework from which experimentally testable predictions can be derived. A first prediction concerns the organization of cortical connectivity during learning. Networks initialized from random connectivity consistently converged to structured connectivity motifs closely resembling those derived n the minimal models, suggesting that these motifs represent general computational solutions rather than artifacts of model construction. We therefore predict that cortical circuits solving the DMC task develop analogous connectivity motifs as learning progresses, even though the underlying biological learning rules differ substantially from backpropagation [28]. Conversely, animals early in learning may rely more strongly on unstructured networks generating mixed selectivity neurons, before structured category-encoding and comparison circuits emerge [12, 15]. Longitudinal recordings combining functional imaging with connectomic reconstruction or targeted perturbation approaches could directly test these predictions by tracking the gradual formation of stable category-encoding populations and their structured projections onto comparison circuits, extending recent studies of learning-dependent representational reorganization [29]. More broadly, the minimal models suggest that increasingly complex behaviors may not require entirely new circuit architectures, but instead arise through the recombination of a limited repertoire of computational motifs implementing operations such as encoding, comparison, temporal coordination and gating. Similar to the extensions of the DMC task considered here, cortical circuits may construct increasingly sophisticated computations by composing these elementary motifs into larger functional architectures [30–34].

Together, our results suggest that heterogeneous and time-varying neuronal responses are not just a byproduct of neural activity, but signatures of an underlying computational architecture that organizes information flow across time. By linking connectivity, neural dynamics, and computation within a unified framework, this work provides a mechanistic account of how structured sequential computations can be generated by structured connectivity in trained circuits.

## IV. ACKNOWLEDGMENTS

A.K. acknowledges funding by the Lise Meitner Excellence Programme of the Max Planck Society. We thank the CCL group members for valuable discussions.

## V. AUTHOR CONTRIBUTIONS

Conceptualization, K.R., A.K; methodology, K.R. A.K.; investigation, K.R.; writing, K.R., A.K.; visualization, K.R.; supervision, A.K.

## VI. METHODS

### A. Modeling network dynamics

We modeled recurrent networks as continuous-time rate-based recurrent neural networks (vanilla RNNs), equivalent to continuous-time Hopfield networks [35]. Each neuron transformed its synaptic input *x*_*i*_ into a firing rate *r*_*i*_ = *ϕ*(*x*) through a nonlinear activation function. The synaptic inputs evolved according to

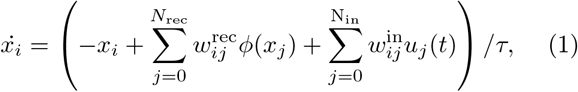

where *w*^rec^ and *w*^in^ denote recurrent and input connection weights, respectively, *u*_*j*_(*t*) are externally controlled inputs and *τ* = 0.1 is the neuronal time constant.

Input populations consisted of direction-selective neurons with von Mises tuning curves,

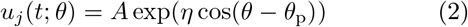

where each input neuron had preferred direction *θ*^*p*^ = 360*/N* ^in^*j*, with *j* ∈ [0, *N* − 1], uniformly distributed over the stimulus space.

Neuronal firing rates were computed using a sigmoidal activation function,

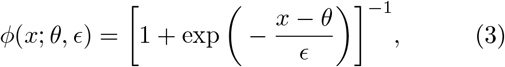

with threshold *σ* = 0.5 and slope parameter *ϵ* = 0.05. This parametrization follows Ref. [36], enabling direct application of its theoretical framework to the construction of the minimal circuits. Output neurons followed the same dynamical equations as recurrent neurons but projected only to themselves. The resulting models can be readily adapted to conventional static readout architectures.

Unless stated otherwise, simulations were performed without stochastic noise. Adding additive white noise preserved the qualitative network dynamics while progressively increasing behavioral errors (Supp. Fig. 3A). The representative trained network shown in the main text was selected for its robustness to dynamical noise. Minimal models were implemented in Julia [37] and integrated using adaptive Runge–Kutta methods provided by *DifferentialEquations*.*jl* [38]. Dynamical systems analyses were performed using *DynamicalSystems*.*jl* [39]. The trained recurrent networks were implemented in Python and simulated using a forward Euler integrator with a sufficiently small integration step, following standard practice in machine learning [40]. The code and parameters for the minimal models is available at https://github.com/KalelR/minimal-cognitive-circuits.

### B. Network training

RNNs were trained using backpropagation techniques [24] with the *Adam* optimizer [41] implemented in PyTorch [40].

The output layer consisted of two neurons representing match (M) and non-match (NM) decisions. During match trials, the target activity of the M neuron was defined by a sigmoid

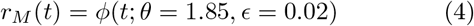

where *ϕ* is the activation function defined in Eq. 3, producing a brief activation during the decoding window [≈1.75, ≈1.9]. The NM neuron remained inactive throughout the trial. Target activities were reversed for non-match trials.

Performance was optimized by minimizing the mean-squared error between network outputs and target activities,

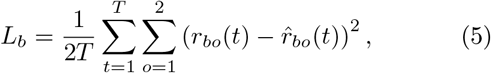

where *r*_*bo*(*t*)_ and 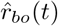 denote the activity and target of output neuron *o* during trial *b*, and *T* is the trial duration. To improve learning robustness, trial losses within each mini-batch were combined using an error-weighted average,

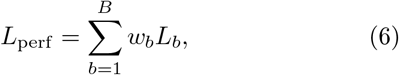

where

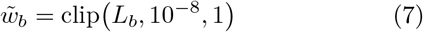

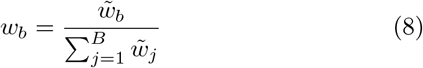

so that

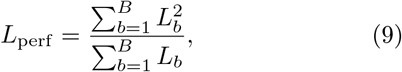

This weighting increases the contribution of trials with larger errors, preventing optimization from focusing primarily on already well-performing trials.

The complete loss term also enforced sparsity constraints on the input, recurrent, and output matrices:

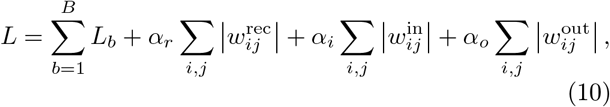

with *α*_*r*_ = 5 *×* 10^−6^, *α*_*i*_ = 10^−6^ and *α*_*o*_ = 3 *×* 10^−6^. To obtain sparse recurrent connectivity, networks were initialized with dense Gaussian-distributed weights and trained iteratively while progressively pruning the weakest 5% of recurrent connections after each training stage. Pruning continued until 12% of the original recurrent connections remained.

The code for the training of the RNNs is available at https://github.com/KalelR/minimal-cognitive-circuits. P. Werbos, Proceedings of the IEEE **78**, 1550 (1990).

