## Supplemental Material for "Structured connectivity for structured sequential computations"

### Supplemental Material for: Structured connectivity for structured computation with categorization and comparison

#### I. COMPARISON OF REPRESENTATION ACROSS MINIMAL NETWORKS

The minimal networks shown in Fig. 2 of the main text encode match or non-match trials using distinct population representations. To characterize these representations, we considered the activity of each neuron as its firing rate averaged over the decoding window ($t \in [1.75, 1.9]$), corresponding to the activity matrices shown in Figs. 2C,F,I.

For the localized network, the population activity associated with each trial type occupies a distinct axis in the neural state space, yielding four mutually orthogonal representations (Fig. S1A). Here, $r_i$ denotes the average activity of neuron $i$. For the hybrid network, AA and BB trials produce the same population activity because both selectively activate the US neuron (Fig. 2D). Consequently, these two trial types are represented along the same axis, whereas AB and BA trials occupy two additional orthogonal axes (Fig. S1B). In this representation, $r_1$, $r_2$, and $r_3$ denote the average activities of the AB, US, and BA neurons, respectively. For the distributed network, we analyzed the representations by projecting the population activity onto two orthogonal vectors,$v_1 = (-1, -1, 1, 1)$ and $v_2 = (1, 1, -1, -1)$. Using the average activities shown in Fig. 2I, and assuming without loss of generality that $r_1 > r_2 > r_3 > r_4 = 0$, the population activity vectors for each trial are AA $= (r_1, r_4, r_2, r_3)$, BB $= (r_4, r_1, r_3, r_2)$, AB $= (r_1, r_3, r_2, r_4)$, BA $= (r_3, r_1, r_4, r_2)$.

Projecting these activities onto the basis $(v_1, v_2)$ yields $x_{AA} = (-r_1 + r_2 + r_3, r_1 - r_2 - r_3)$, $x_{BB} = (-r_1 + r_2 + r_3, r_1 - r_2 - r_3)$, $x_{AB} = (-r_1 - r_3 + r_2, r_1 + r_3 - r_2)$ and $x_{BA} = (-r_1 - r_3 + r_2, r_1 + r_3 - r_2)$. Thus, match ($x_{AA} = x_{BB}$), project to the same point in this representation, as do non-match trials ($x_{AB} = x_{BA}$). Fig. S1C, illustrates this projection using values similar to those of the minimal networks ($r_1 = 1$, $r_2 = 0.8$, $r_3 = 0.5$).

#### II. LOCALIZED INSTEAD OF DISTRIBUTED DECODING FOR IDENTICAL NETWORK ACTIVITY

The distributed minimal network shown in the main text (Fig. 2G-I) decodes the comparison through purely excitatory connections, requiring the combined activity of multiple neurons to distinguish trial types. However, the population activity in the distributed network (Fig. 2I) can also be interpreted as the complement of the localized representation (Fig. 2C), in which the inactive neuron identifies the trial type. This representation can therefore be decoded using inhibitory rather than excitatory readout connections.

To demonstrate this, we constructed a minimal network with the same population activity as the distributed minimal network but a localized decoding scheme (Fig. S2). In this network, all comparison neurons (C1, C2, R, and B) inhibit the output layer, with A and B inhibiting the match (M) output neuron and C1 and C2 inhibiting the non-match (NM) output neuron. An additional non-selective neuron (US) provides tonic excitation to both output neurons. Because this excitation would otherwise activate the outputs prematurely, a sample neuron was included to gate the output layer during the comparison period. This example illustrates that the computation performed by the network depends not only on the population representation but also on the structure of the downstream decoder.

#### III. EXAMPLES OF ADDITIONAL TRAINED RNNS

We trained networks using multiple random initializations and a range of training hyperparameters, including the learning rate and sparsity regularization. Some parameter combinations failed to converge to a functional solution. From each training batch, we selected the network with the highest robustness to additive noise for further analysis. Figure S3B-F shows an additional trained network (seed 7 in Figure S3A) obtained from a different initialization. This network achieved the highest performance under additive white noise within its training batch. Panels B-F reproduce the analyses shown in Fig. 4 of the main text. As in the primary example, training gave rise to category-selective neurons, mixed-selectivity neurons, and neurons with no apparent task selectivity. Mixed-selectivity neurons included both localized responses, active during a single trial type, and distributed responses, active during multiple trial types.

*
†

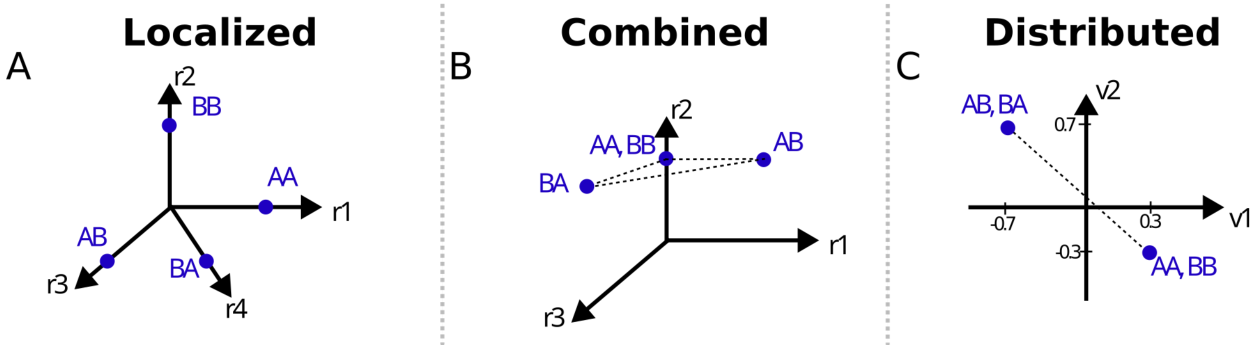


Figure S1. Representation of average activity during the decoding period across trials for the minimal networks shown in Fig.2).

#### IV. TRACES OF ALL NEURONS IN THE TRAINED NETWORK

In the main text, we presented representative traces from the trained network (Fig.4). Fig. S4 contains all of the neuronal activity traces. The dynamics of the network is relatively sparse, with many neurons remaining largely inactive as a consequence of the sparsity constraint imposed during training. It is important to note here that two neurons (nodes 13 and 44) exhibit approximately localized mixed selectivity for category BB. These neurons were omitted from the main figure because they project only weakly to the output layer and therefore make a negligible contribution to task performance. More generally however, this illustrates that the trained recurrent networks need not to exploit all available neurons efficiently and can contain functionally redundant units.

#### V. POPULATION DYNAMICS AND FIXED POINT ANALYSIS OF THE MINIMAL AND TRAINED NETWORKS

To characterize the network dynamics, we identified stable fixed points for the baseline system (no stimulus) and for representative stimuli from each category, and projected the neural trajectories onto the first three principal components (Fig. S5). The trajectories corroborate the circuit analysis presented in the main text while providing a population-level view of the computation. The localized minimal network (Fig. S5B) contains two stimulus-dependent fixed points during the sample epoch, corresponding to the two possible category contexts, and four fixed points during the test epoch, one for each sample-test category combination. During the sample period, the trajectory follows a transient toward a stimulus-dependent fixed point without reaching it, preventing premature readout. During the delay, the trajectory converges to a baseline fixed point encoding the sample category, and during the test it approaches the decision-specific fixed point before returning to the baseline attractor. The trained and hybrid networks exhibit qualitatively similar dynamics (Fig. S5A,C). Although the distributed network follows the same overall organization, it contains an additional stimulus-dependent fixed point arising from the sample-selective population (Fig. S5D). These results show that the computation is organized by transitions between transient dynamics and stable attractors. The resulting neural trajectories must be interpreted together with the downstream readout, as the network output depends on both the population state and the connectivity to the output layer.

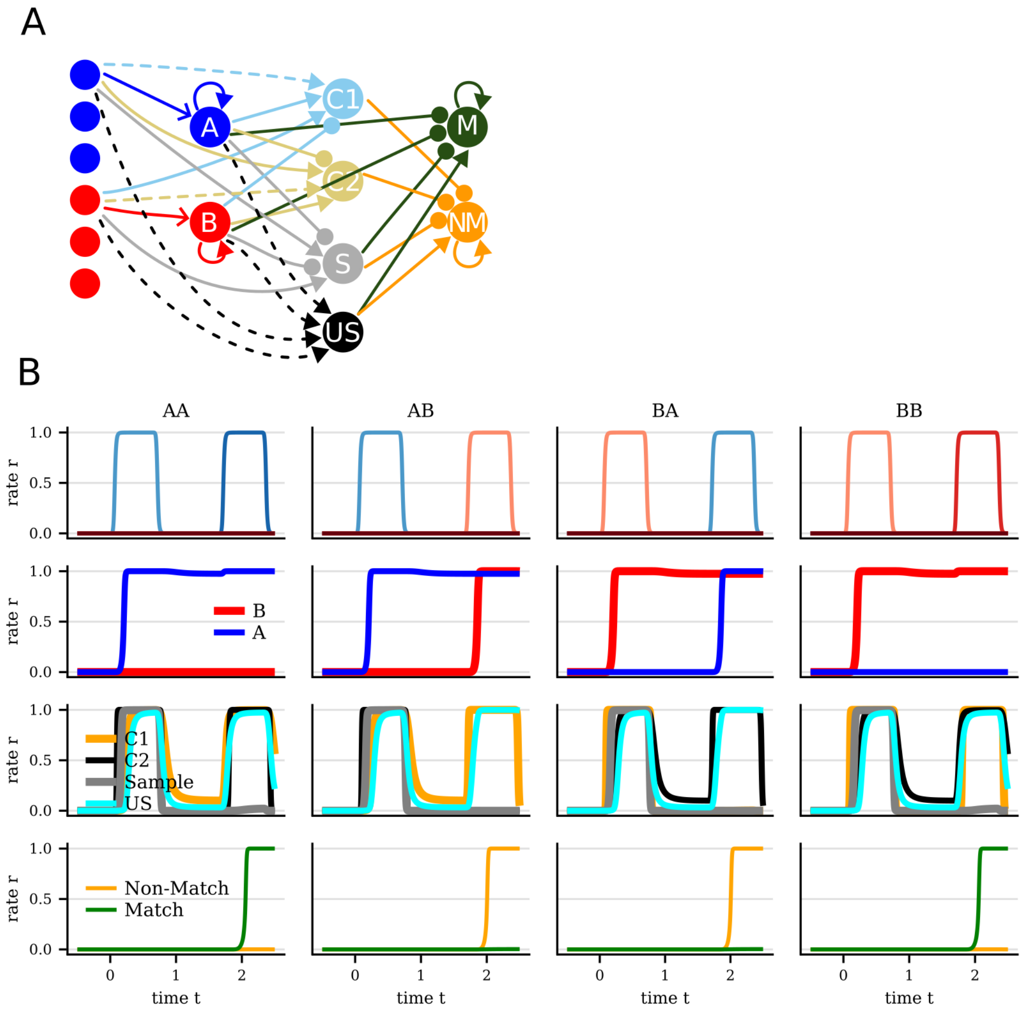


Figure S2. **Distributed population representation with localized decoding.** The network reproduces the population activity of the distributed minimal network shown in Fig. 2 while using a localized decoding scheme. A non-selective (US) neuron provides tonic excitation to the output layer, whereas comparison neurons inhibit the appropriate output neurons (A and B inhibit the M output; C1 and C2 inhibit the NM output neurons). Consequently, trial identity is decoded from the inactive comparison neuron rather than from the collective excitatory activity of multiple neurons.

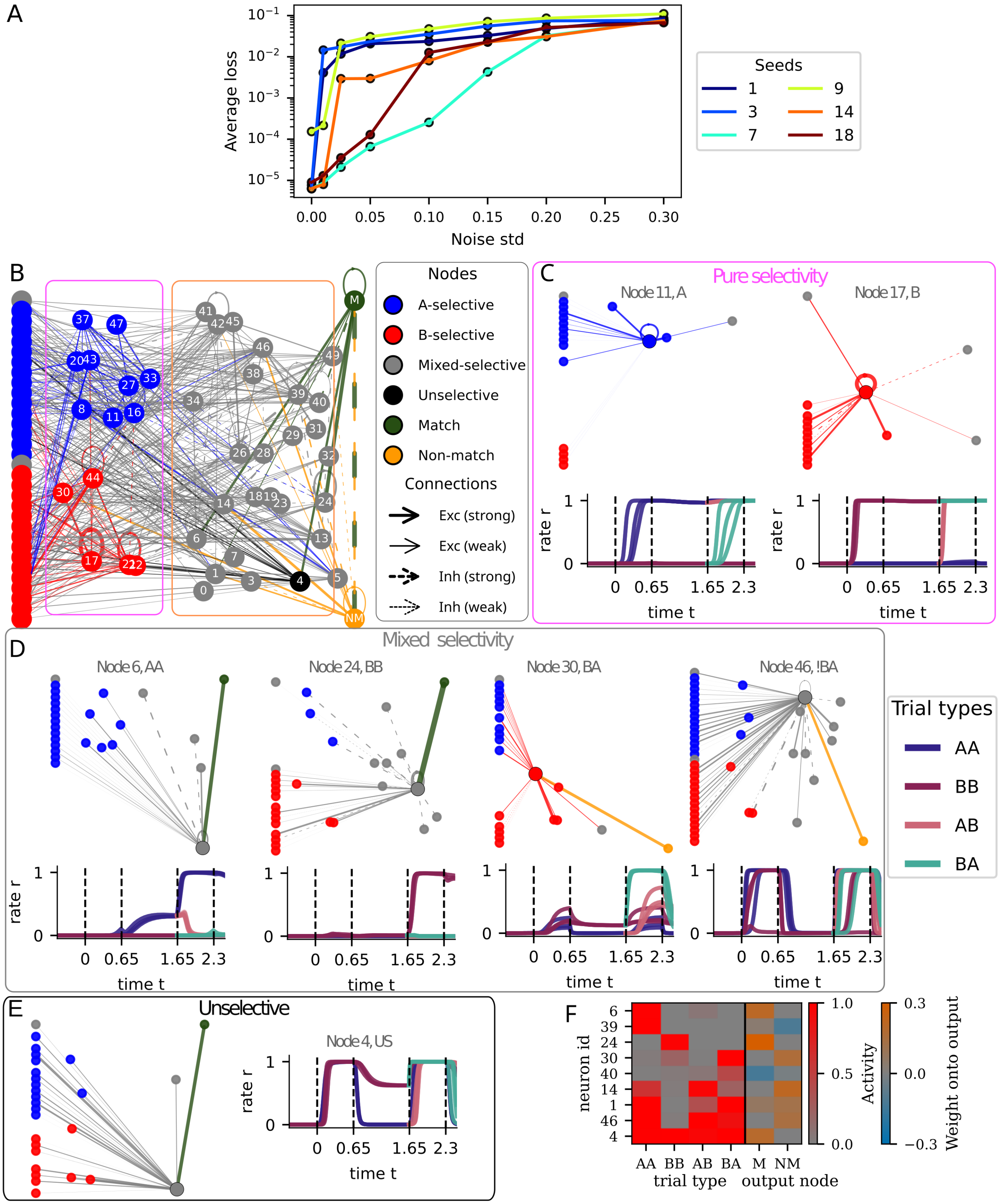


Figure S3. **Connectivity and selectivity organization in an additional trained network example.** (A) Average loss of different trained network examples with increasing noise intensity. (B) Connectivity structure of a representative trained RNN obtained from a random initialization (seed 7 in (A)). Nodes are colored and positioned according to their selectivity class, identified from their activity profiles, and edge widths indicate the magnitude of synaptic weights. For visualization, weak connections and nodes lacking strong connections are omitted. (C–E) Connectivity and activity profiles of representative neurons from the trained network. Top, incoming and outgoing connections of selected neurons; bottom, neuronal responses across all stimulus combinations, grouped by trial type. (F) Activity and output connectivity of neurons with the strongest influence on decision generation. Left, mean firing rates during the decoding period for each input combination within a trial type (gray-to-red scale). Right, corresponding output weights onto match and non-match decision neurons (blue-to-brown scale).

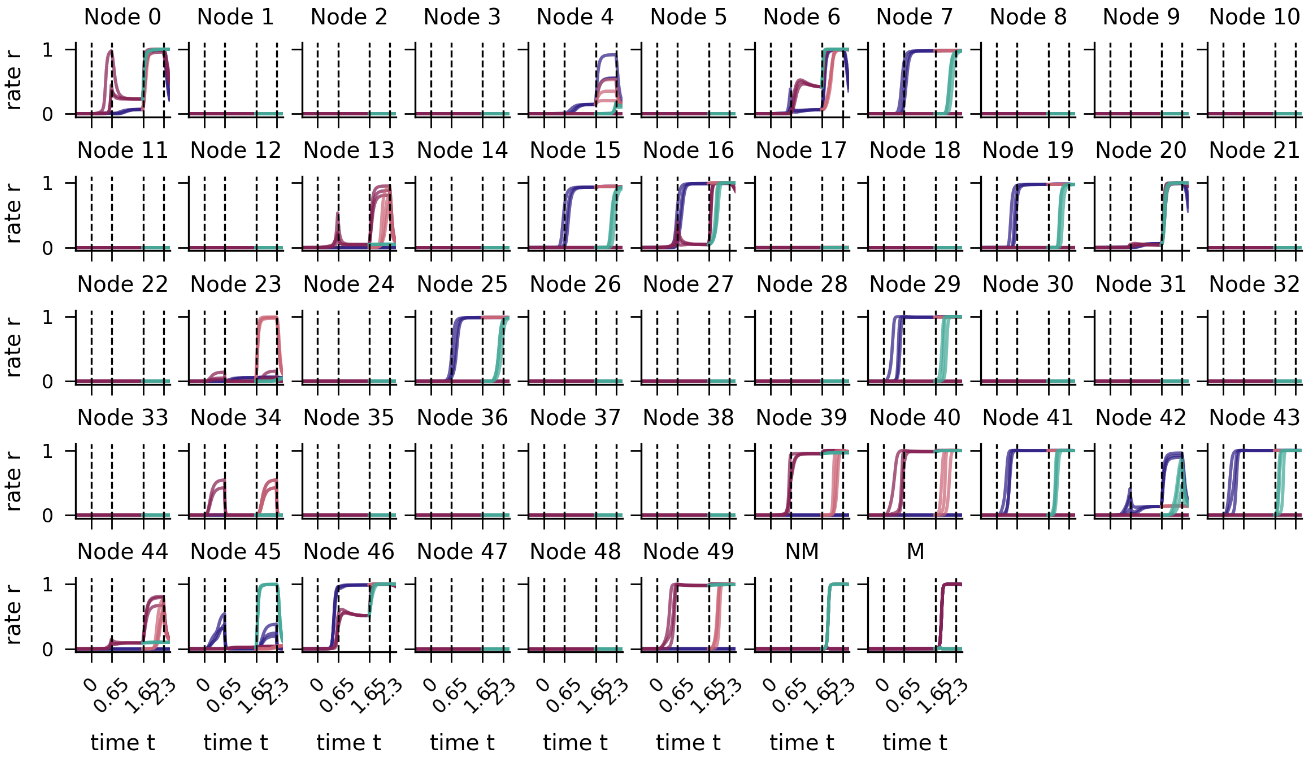


Figure S4. **Neuronal activity traces.** Activity traces for all neurons in the sparse trained network shown in Fig. 4. Colors follow the selectivity classification used throughout the paper. The last two panels show the activities of the non-match (NM) and match (M) output neurons.

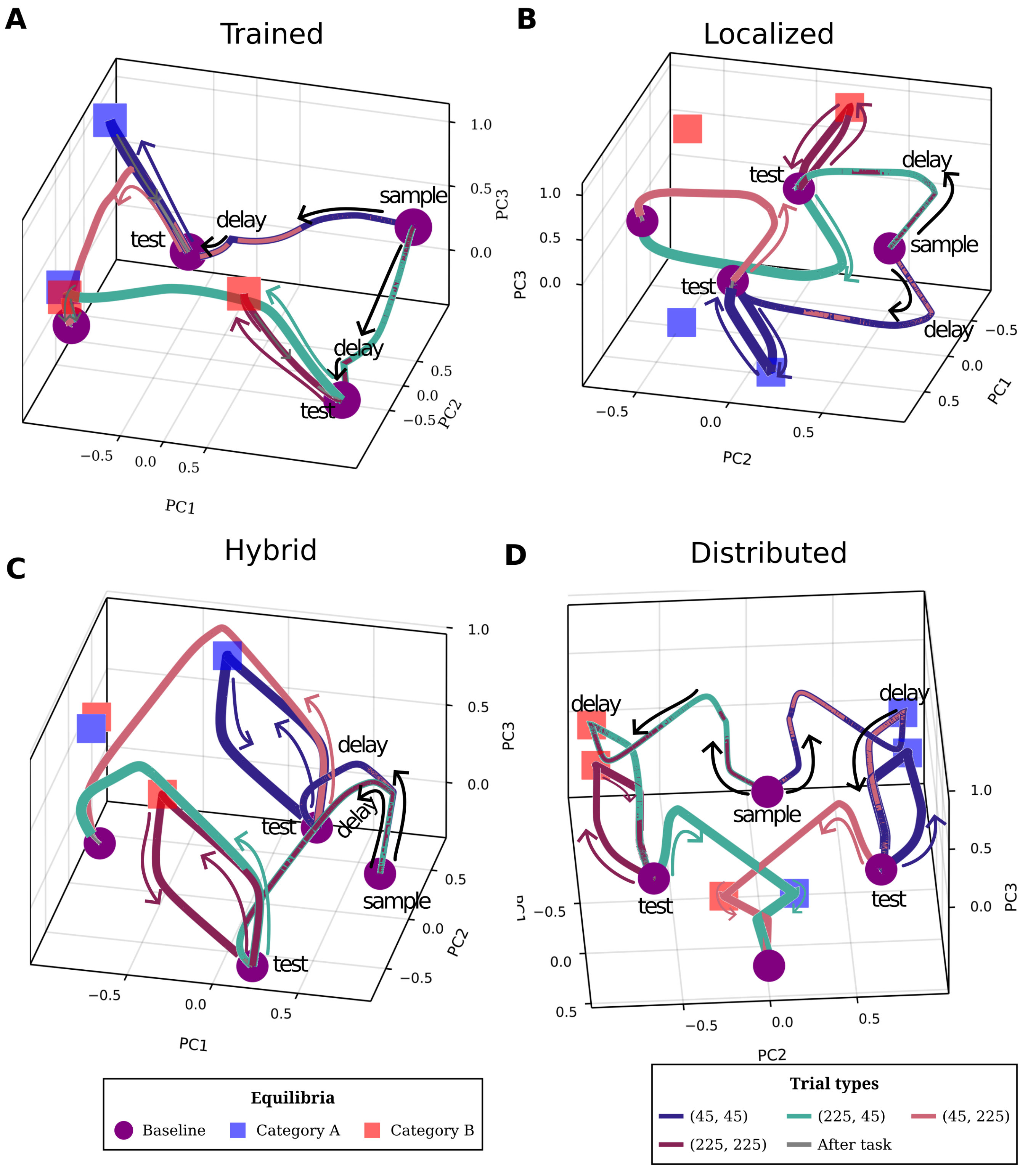


Figure S5. **Population dynamics of the trained and minimal networks**. (A) Evolution of the activity trajectory of the trained network shown in Fig.4, projected onto the first three principal components. The start of the corresponding trial epoch is denoted, with a single trajectory per trial type. Starting in the sample epoch, trajectories separate out in the projected state space, switching between transients to fixed points as the stimuli are presented. (B-D) analogous to (A), but for the minimal networks, shown in Fig.2A, D, G.
